# Myc-induced cell mixing is required for competitive tissue invasion and destruction

**DOI:** 10.1101/020313

**Authors:** Romain Levayer, Barbara Hauert, Eduardo Moreno

## Abstract

Cell-cell intercalation is used in several developmental processes to shape the normal body plan^1^. There is no clear evidence that intercalation is involved in pathologies. Here, we use the proto-oncogene Myc to study a process analogous to early phase of tumour expansion: Myc-induced cell competition^2-7^. Cell competition is a conserved mechanism^5,6,8,9^ driving the elimination of slow proliferating cells (so called losers) by faster proliferating neighbours (so called winners) through apoptosis^10^ and is important to prevent developmental malformations and maintain tissue fitness^11^. Using long term live imaging of Myc-driven competition in the *Drosophila* pupal notum and in the wing imaginal disc, we show that the probability of elimination of loser cells correlates with the surface of contact shared with winners. As such, modifying loser/winner interface morphology can modulate the strength of competition. We further show that elimination of loser clones requires winner/loser cell mixing through cell-cell intercalation. Cell mixing is driven by differential growth and the high tension at winner-winner interfaces relative to winner-loser and loser-loser interfaces, which leads to a preferential stabilisation of winner-loser contacts and reduction of clone compactness over time. Differences in tension are generated by a relative difference of junctional F-actin levels between loser and winner junctions, induced by differential levels of the phosphatidylinositol PIP3. Our results establish the first link between cell-cell intercalation induced by a proto-oncogene and how it promotes invasiveness and destruction of healthy tissues.

---

To analyse quantitatively loser cell elimination, we performed long term live imaging of clones showing a relative decrease of the proto-oncogene Myc in the *Drosophila* pupal notum (**Fig. 1a,b, Video S1**), a condition known to induce cell competition in the wing disc^3,4^. Every loser cell delamination was counted over 10h and we calculated the cell elimination probability for a given surface of contact shared with winner cells (**Fig. 1c,d** and Methods). We observed a significant increase of the proportion of delamination with winner/loser shared contact, while this proportion remained constant for control clones (**Video S2**, **Fig. 1d**). The same correlation was observed in *ex-vivo* culture of larval wing disc (**EDFig.1**, **Video S3**). Cell delamination in the notum was apoptosis dependent (**Video S4**, *UAS-diap1*, 1/922 cells delaminated, 4 nota) and expression of *flower*^*lose*^ (*fwe*), a competition-specific marker for loser fate^12^, was necessary and sufficient to drive contact dependent delamination (**Fig. 1d**, **Video S5-S6**). Moreover we confirmed^11,12^ that contact-dependent death is based on the computation of relative differences of *fwe*^*lose*^ between loser cells and their neighbours (**Ext. Fig 2a-e**). Thus, cell delamination in the notum recapitulates features of cell-competition^4,12^.

**Figure 1:**
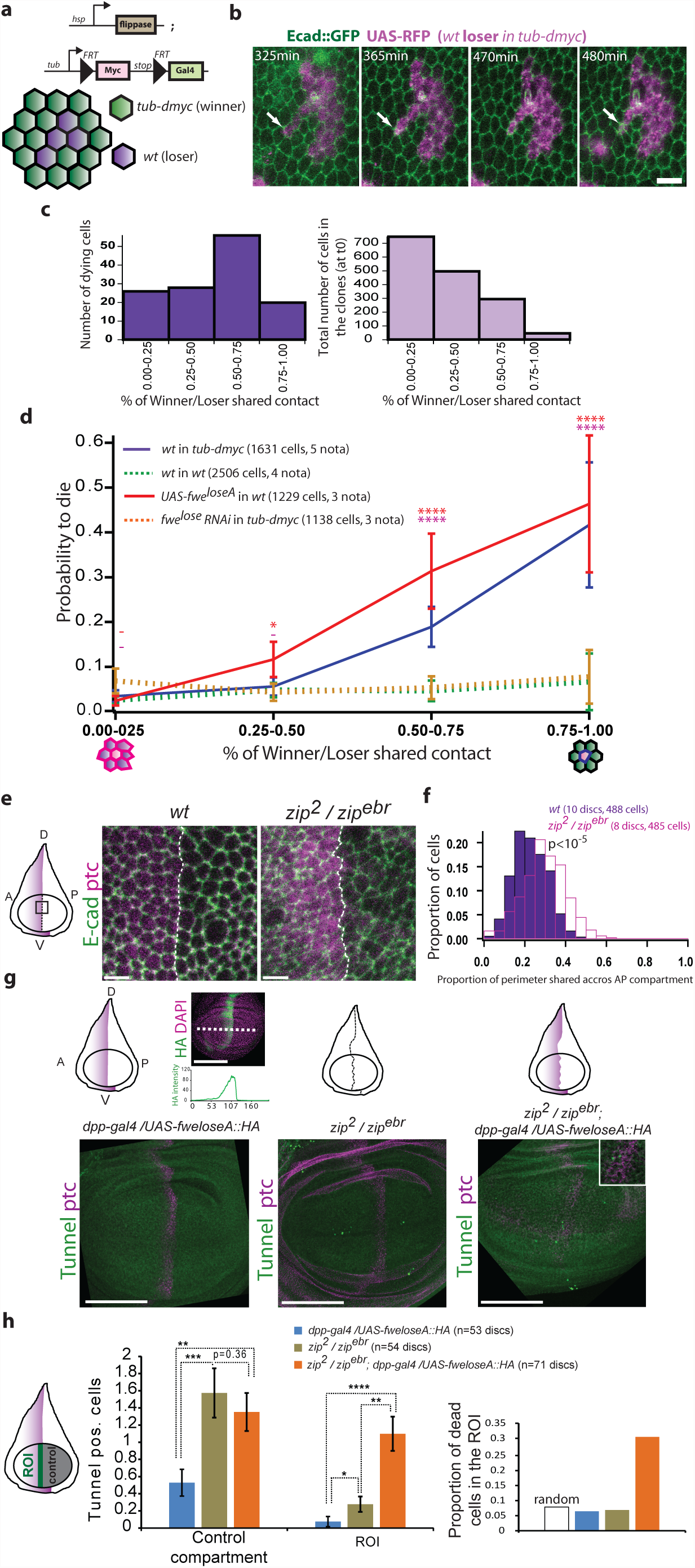
Loser elimination correlates with the surface shared with winners. **a,** Description of the supercompetition assay^4^. **b,** Snapshots of Video S1 showing loser cell delamination (white arrowheads) in the pupal notum (a single layer epithelium). Scale bar=10μm. **c,** Distribution of the proportion of junctional perimeter of loser cells shared with winner cells 1 hour prior to delamination (top) and in all the loser cells at t0 (bottom). **d,** Probability of loser cell elimination for a given surface of contact shared with winners. Statistical tests are Fisher-exact tests performed with *wt* in *wt*. Error bars are 95% confidence intervals. **-**: p>0.05, *:p<0.05, **:p<10^−2^, ***: p<10^−3^, ****: p<10^−4^. **e,** Adherens junctions (E-cad) of cells at the wing disc AP compartment boundary (white dotted line, localized with Patched (ptc)) in *wt* and upon downregulation of MyosinII heavy chain (*zip*^*2*^*/zip*^*ebr*^). (A: Anterior, P: Posterior, D: Dorsal, V: Ventral). Scale bars = 5μm. **f**, Distribution of the proportion of perimeter shared with cells across the compartment boundary (p, Mann-Whitney test). **g**, Z-projection of a wing disc overexpressing *fwe*^*loseA*^:*:HA* in the anterior compartment (left), a *zip*^*2*^*/zip*^*ebr*^ disc (middle) and a disc combining both (right, inset=tunnel positive cells at the AP boundary). Top schematic: purple=*fwe*^*loseA*^:*:HA,* black line=AP compartment boundary. Note that *fwe*^*loseA*^:*:HA* is expressed in a graded way in the anterior compartment (top left). Scale bars = 100μm. **h**, Average number of Tunnel positive cells in the posterior compartment (control) and in vicinity of the AP boundary (ROI, top scheme). Bottom: ratio of average number of dead cells in the ROI over the total number of dead cells in the wing pouch. The white box is the expected ratio for a random distribution (ROI surface/total wing pouch surface). p=Mann-Whitney test.

This suggested that winner/loser interface morphology could modulate the probability to eliminate loser clones. Using the wing imaginal disc, we reduced winner/loser contact by inducing adhesion or tension-dependent cell sorting^13^ (**Fig. 2d**) and observed a significant reduction of loser clone elimination (**EDFig.3a-c, EDFig.2f,g**). This rescue was not driven by a cell autonomous effect of E-cadherin (E-cad) or active MRLC (MyosinII Regulatory Light Chain) on growth, death or cell fitness (**EDFig.3d-f**, **Videos S7-S8**) but rather by a general diminution of winner/loser contact. Competition is ineffective across the Antero-Posterior (AP) compartment boundary^14^, a frontier that prevents cell mixing through high line tension^15^. Accordingly, there was no death increase at the AP boundary in wing discs overexpressing *fwe*^*loseA*^ in the anterior compartment (**Fig. 1 g,h**). However, reducing tension by reducing MyosinII Heavy Chain levels was sufficient to increase the shared surface of contact between cells of the A and P compartment (**Fig. 1 e,f**, p<10^−5^), and elicited *fwe*^*lose*^ death induction at the boundary (**Fig. 1 g,h**). Altogether, we concluded that the reduction of surface of contact between winners and losers is sufficient to block competition and this explains how compartment boundaries prevent competition.

**Figure 2:**
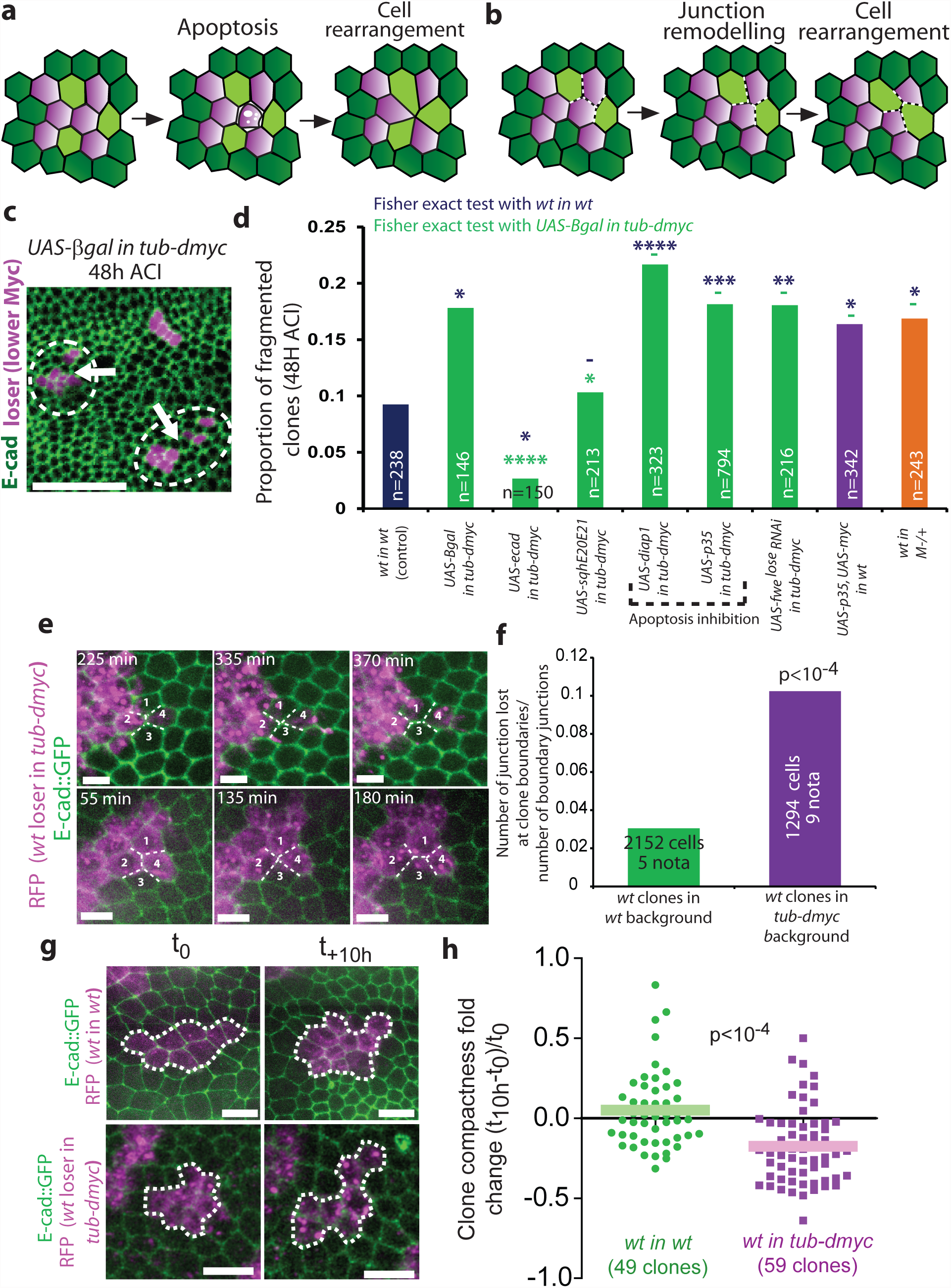
Winner/loser mixing is induced by junction remodelling and cell-cell intercalation. **a,** Clone fragmentation induced by cell death and the subsequent cell rearrangement **b,** Clone fragmentation induced by cell-cell intercalation **c,** Top left, supercompetition assay in the wing disc 48h ACI (loser cells, purple). White arrowheads=fragmented clones. Scale bar = 25μm. **d,** Proportion of fragmented clones (images in EDFig.4c). n=number of clones. (test: Fisher-exact tests, **-**: p>0.05, *:p<0.05, **:p<10^−2^, ***: p<10^−3^, ****: p<10^−4^). **e,** Junction remodelling events in loser clones in the pupal notum 48h ACI leading to clone splitting (top) or persistent disappearance of a loser-loser junction (bottom). Scale bars = 5μm. **f,** Proportion of RFP/RFP junction at clone boundaries disappearing through remodelling after 12h in the pupal notum (p: Fisher-exact test). **g**, *wt* in *wt* clone and *wt* in *tub-dmyc* clone in the notum at time 0 and 10 hours later (48h ACI, 20h after pupae formation). Scale bars = 5μm. **h**, Evolution of single clone compactness over time in the notum ((t10h-t0)/t0, see Methods). One dot = one clone, bars are averages. p: Mann-Whitney test.

Loser clones have been reported to fragment more often than control clones^14,16^, while winner clones show convoluted morphology^14,17^, suggesting that winner-loser mixing is increased during competition. This could affect the outcome of cell competition by increasing the surface shared between losers and winners. We used clone splitting as a readout for loser-winner mixing. Two non-exclusive mechanisms can drive clone splitting: cell death followed by junction rearrangement (**Fig. 2a**) or junction remodelling and cell-cell intercalation independently of death (**Fig. 2b**). To assess the contribution of each phenomenon, we systematically counted the proportion of clones fragmented 48 hours after clone induction (**Fig. 2c**, see Methods and **EDFig.4a,b**). We observed a two fold increase of the frequency of split clones in losers (*wt in tub-dmyc*) versus *wt in wt* controls (**Fig. 2d, EDFig.4c**). Overexpressing E-cad or active MyoII was sufficient to prevent loser clone splitting while blocking apoptosis or blocking loser fate by silencing *fwe*^*lose*^ did not reduce splitting (**Fig. 2d, EDFig.4c**). Finally, the proportion of split clones was also increased for winner clones either during Myc-driven competition (*UAS-myc, UAS-p35*, **Fig. 2d, EDFig.4c**) or during Minute-dependent competition^2^ (*wt* clones in *M*^−^*/+* background, **Fig. 2d, EDFig.4c**). Altogether, this suggested that winner/loser mixing is increased independently of loser cell death or clone size (**EDFig.4d**) by a factor upstream of *fwe,* and could be driven by cell-cell intercalation. Accordingly, junction remodelling events leading to disappearance of a loser-loser junction were three times more frequent at loser clone boundaries compared to control clone boundaries in the pupal notum (**Fig. 2e,f**, p<10^−4^, **Video S9**). The rate of junction remodelling was higher in loser-loser junctions and in winner-winner junctions compared to winner-loser junctions (**EDFig.5a,b**). The preferential stabilisation of winner-loser interfaces should increase the surface of contact between winner and loser cells over time. Accordingly, loser clone compactness in the notum was decreasing over time while it remains constant on average for *wt* clones in *wt* background (**Fig. 2g,h**, p<10^−4^). Similarly, the compactness of clones in the notum was also decreasing over time for conditions showing high frequency of clone splitting in the wing disc, while clone compactness remained constant for conditions rescuing clone splitting (compare **Fig. 2d** with **EDFig.5 d,e**, **Videos S1-S11**). Altogether, we concluded that both Minute and Myc-dependent competition increase loser/winner mixing through cell-cell intercalation.

We then asked what could modulate the rate of junction remodelling during competition. The rate of junction remodelling can be cell-autonomously increased by Myc (**EDFig.5c**, **Video S12**). Interestingly, downregulation of the tumour suppressor PTEN is also sufficient to increase the rate of junction remodelling^18^ through the upregulation of the phosphatidylinositol PIP3. We reasoned that differences in PIP3 levels could also modulate junction remodelling during competition. Using a live reporter of PIP3 which can detect modulations of PIP3 in the notum (**EDFig.6a,b**), we observed a significant increase of PIP3 in the apico-lateral membrane of *tub-dmyc/tub-dmyc* interfaces compared to *wt/wt* and *wt/tub-dmyc* interfaces (**Fig. 3 a,b**). Moreover, increasing/reducing Myc levels in a full compartment of the wing disc was sufficient to increase/decrease the levels of phospho-Akt (a downstream target of PIP3^19^, **Fig. 3c**) while *fwe*^*loseA*^ overexpression had no effect (**EDFig.6c**). Similarly, levels of phospho-Akt were relatively higher in *wt* clones compared to the surrounding *M*^−^*/+* cells (**EDFig.6d**). Thus differences in PIP3 levels might be responsible for winner-loser mixing. Accordingly, reducing PIP3 levels by overexpressing a PI3K-DN (PI3 Kinase Dominant Negative) or increasing PIP3 levels by knocking down PTEN (*UAS-pten RNAi*) were both sufficient to induce a high proportion of fragmented clones (**Fig. 3d,e**) and to reduce clone compactness over time in the notum (**EDFig.5d,e**, **Video S13, S14**), while increasing PIP3 in loser clones was sufficient to prevent cell mixing (**Fig. 3d,e**). Moreover, abolishing winner-loser PIP3 differences through larval starvation (^20^, **EDFig.6e-g**) prevented loser clone fragmentation (**EDFig.6h,i**), the reduction of clone compactness over time in the notum (**EDFig.5d,e**, **Video S15**) and could rescue *wt* clone elimination in *tub-dmyc* background (**EDFig.6 j,k**). We therefore concluded that differences in PIP3 levels are necessary and sufficient for loser-winner mixing and required for loser cell elimination.

**Figure 3:**
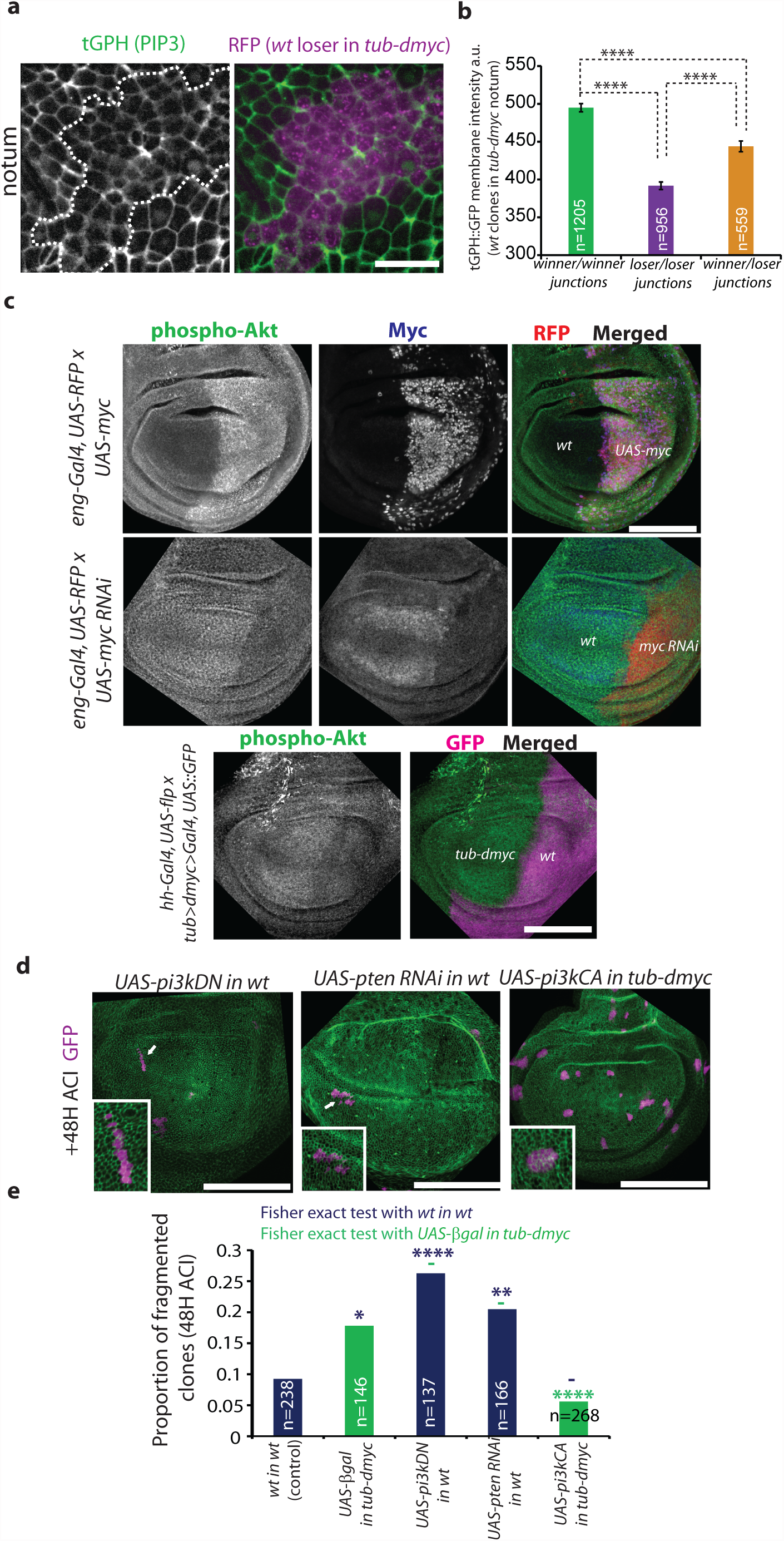
Differences in PIP3 induce loser-winner mixing. **a,** Z-projection of tGPH (PIP3) around the apico-lateral domain of loser clones (purple) in the pupal notum 48h ACI (White dashed lines=clone boundaries). Scale bars = 10μm. **b,** Mean membrane intensity of tGPH. n=number of junctions., ****: p<10^−4^, Mann-Whitney test. Error bars=s.e.m. **c,** Z-projections of phospho-Akt (green), RFP (red) and Myc (blue) in wing discs upon modulation of Myc levels in the posterior compartment. Scale bars=100μm. Note that intensities are not comparable between discs. (n=10 discs for each). **d**, Wing discs 48h ACI with clones with low PIP3 (UAS-PI3KDN, low PIP3) or high PIP3 (*UAS-pten RNAi*) or loser cells (supercompetition assay) overexpressing a constitutive active form of PI3K (PI3K-CA). Insets show representative clones. White arrows=fragmented clones. Scale bars=100μm. **e,** Proportion of fragmented clones. n=number of clones. p=Fisher-exact tests, **-**: p>0.05, *:p<0.05, **:p<10^−2^, ***: p<10^−3^, ****: p<10^−4^. *wt* in *wt* and *wt* in *tub-dmyc* come from Fig. 2d.

We then asked which downstream effectors of PIP3 could affect junction stability. A relative growth decrease can generate mechanical stress^21,22^ that can be released by cell-cell intercalation. Accordingly, growth reduction through Akt downregulation is sufficient to increase clone splitting (**EDFig.7 a,b**) and could contribute to loser clone splitting. However, Akt is not sufficient to explain winner-loser mixing as unlike PIP3, increasing Akt had no effect on clone splitting (**EDFig.7 a,b**). PIP3 could also modulate junction remodelling through its effect on cytoskeleton^23^ and the modulation of intercellular adhesion or tension^1^. We could not detect obvious modifications of E-cad, MRLC or Dachs (another regulator of tension^24^) in loser cells (**EDFig.7c-f**). However, we observed a significant reduction of F-actin levels and a reduction of actin turnover/polymerisation rate in loser-loser and loser-winner junctions in the notum (**Fig. 4a,b**, p<10^−5^, **EDFig.8, Video S16-S17**). Similarly, modifying Myc levels in a full wing disc compartment was sufficient to modify actin levels (**Fig. 4c**), and F-actin levels were higher in *wt* clones compared to *M-/+* cells (**EDFig.9a**). This prompted us to test the role of actin organisation in winner/loser mixing. Downregulating the formin Diaphanous (a filamentous actin polymerisation factor^13^) by RNAi or by using a hypomorphic mutant was sufficient to obtain a high proportion of fragmented clones (**Fig.4d**, **EDFig.9b**, 28%, p<10^−4^, **EDFig.9c**, 39% fragmented) and to reduce clone compactness over time (**EDFig.5d,e**, **Video S18**), while overexpressing Dia in loser clones prevented clone splitting (*UAS-dia::GFP*, **Fig.4d**, **EDFig.9b**) and compactness reduction (**EDFig.5d,e**, **Video S19**). This effect was specific of Dia as modulating Arp2/3 complex (a regulator of dendritic actin network^13^) had no effect on clone splitting (**Fig.4d**, **EDFig.9b**). Thus, impaired filamentous actin organisation was necessary and sufficient to drive loser-winner mixing. These actin defects were driven by the differences in PIP3 levels between losers and winners (see **EDFig.10**). Thus Diaphanous could be an important regulator of competition through its effect on cell mixing. Overexpression of Dia was indeed sufficient to significantly reduce loser clone elimination (**EDFig.9d**) without affecting Hippo/YAP-TAZ pathway^25^ (**EDFig.9e**).

**Figure 4:**
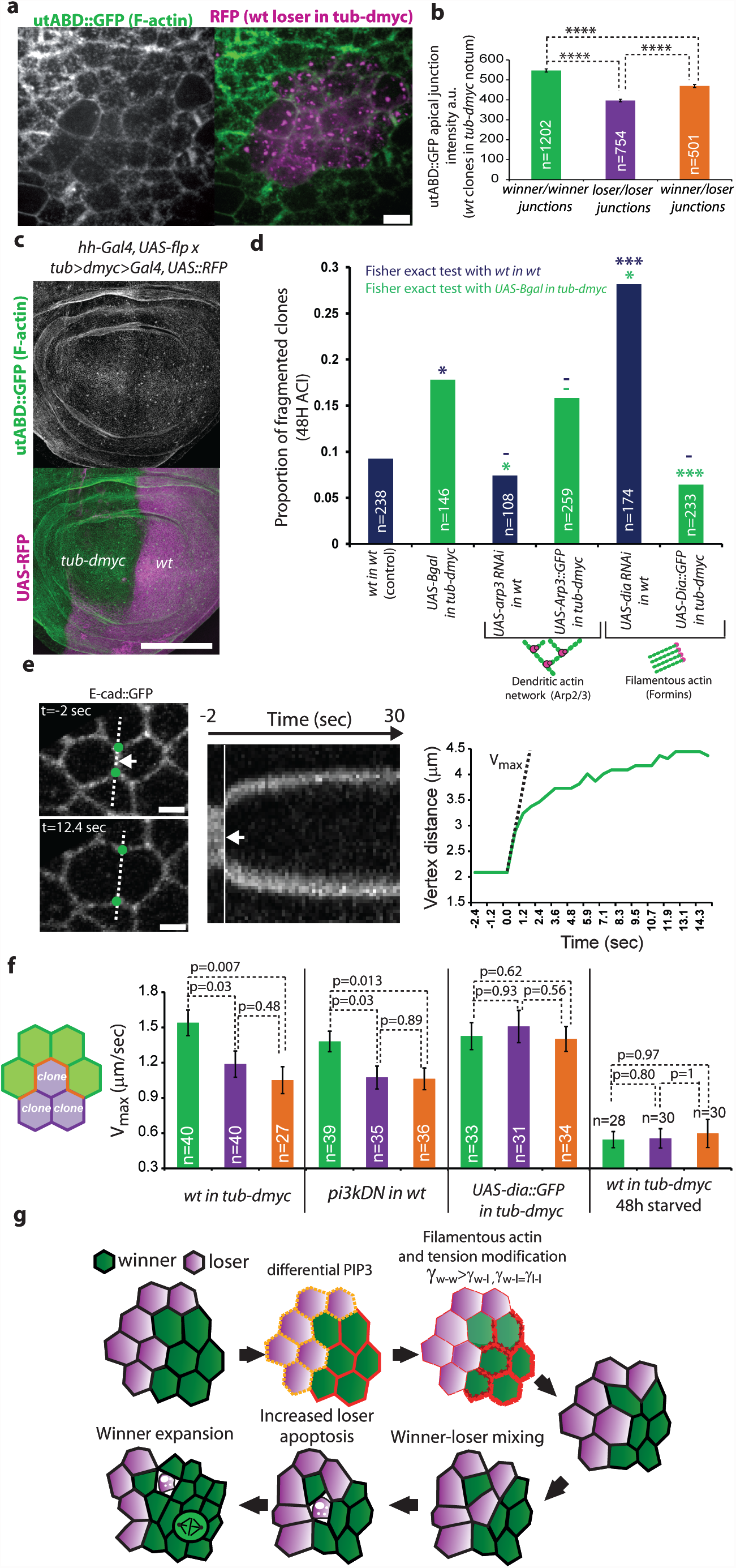
Filamentous actin and tension modulation are responsible for winner-loser mixing. **a,** Z-projection around junction plane in the pupal notum 48h ACI of utABD::GFP (utrophin Actin Binding Domain, F-actin) during supercompetition (purple cells: losers). Scale bar=5μm. **b,** Mean junctional utABD::GFP intensity. a.u.: arbitrary units. n=number of junctions. Error bars are s.e.m.. p= Mann-Whitney tests. **c,** Z-projections of utABD::GFP (green) and RFP (magenta) in a wing disc after removal of one additional copy of Myc in the posterior compartment (16/16 discs). Scale bar=100μm. **d,** Proportion of fragmented clones upon perturbation of actin. Discs are shown in EDFig.9b. n=number of clones. **-**: p>0.05, *:p<0.05, **:p<10^−2^, ***: p<10^−3^, ****: p<10^−4^. Fisher-exact tests. *wt* in *wt* and *wt* in *tub-dmyc* come from Fig. 2d. **e,** Junction laser nanoablation in the notum before ablation (top) and 12.4sec after ablation (bottom). White arrow=ablation point, green points= vertices. Right: kymograph showing the evolution of distance between vertices over time and vertices distance ploted over time. The Vmax (maximum speed of relaxation following ablation) is calculated for each curve and used to assess interfacial tension (see Methods). **f**, Mean Vmax after junction ablation (green: winner-winner, purple: loser-loser, orange: winner-loser). n=number of junctions, p-values are Mann-Whitney tests. **g**, Model of the winner-loser mixing process (see Supplementary discussion). γ =interfacial tension, w=winner, l=loser, red arrows=increased interfacial tension, red lines=F-actin.

Filamentous actin has been associated with tension regulation^13^. We therefore asked whether junction tension was modified in winner and loser junctions. The maximum speed of relaxation of junction following laser nanoablation (which is proportional to tension^26^) was significantly reduced in loser-loser and winner-loser junctions compared to winner-winner junctions (**Fig. 4e,f, Video S20**). This distribution of tension has been proposed to promote cell mixing^27^. Accordingly, decreasing PIP3 in clones reduced tension both in *low-PIP3/low-PIP3* and *low-PIP3/normal-PIP3* junctions, while overexpressing Diaphanous in loser clones or starvation were both sufficient to abolish differences in tension (**Fig. 4f, Videos S21-S23**), in agreement with their effect on winner/loser mixing and the distribution of F-actin. Thus the lower tension at winner/loser and loser/loser junctions is responsible for winner/loser mixing. Altogether, we concluded that the relative PIP3 decrease in losers increases winner-loser mixing through Akt-dependent differential growth and the modulation of tension through F-actin downregulation in winner/loser and loser/loser junctions (**Fig. 4g**).

Several modes of tissue invasion by cancer cells have been described^28^, most of them relying on the departure of the tumour cells from the epithelial layer^29^. This study suggests that some oncogenes may also drive tissue destruction and invasion by inducing ectopic cell intercalation between cancerous and healthy cells, and subsequent healthy cell elimination. Myc dependent invasion could be enhanced by other mutations further promoting intercalation (such as *PTEN*). Stiffness is increased in many tumours^30^, suggesting that healthy cells/cancer cells mixing by intercalation might be a general process.

## Acknowledgments

We would like to thank members of the Moreno lab for critical reading of this manuscript. We also would like to thank M. Bergen for his help on E-cad FRAP data collection. We are also very grateful to Y. Bellaïche, O. Baumann, J. Grossahns, H. Jasper, T. Lecuit, L. Legoff, G. Morata, H. Stocker, R. Sousa-Nunes, the Bloomington stock center and the DSHB for sharing stocks and reagents, to B. Aigouy for the Packing analyser software and the Center for Microscopy and Image Analysis (University of Zurich) for sharing equipment.

R.L. was supported by an EMBO long term fellowship (ALTF 366-2012) and a Human Frontier post-doctoral fellowship (LT000178/2013). Work in our laboratory is funded by the European Research Council, Swiss National Science Foundation, Josef Steiner Cancer Research Foundation, and the Swiss Cancer League.

## Author contribution

R.L. and E.M. designed the experiments. R.L. performed and analysed the experiments. B.H. generated *fwe* KO and *fwe*^*loseA*^:*:mcherry* knock in. R.L. and E.M. wrote the manuscript.

## Authors information

Reprints and permissions information is available at www.nature.com/reprints. The authors declare no competing interest. Correspondence and requests for materials should be addressed to.

## Supplementary discussion

### Proposed model for winner-loser mixing

Two forces contribute to winner-loser mixing:

1. The relative decrease of growth of loser clones. Relative growth decrease leads to clone stretching^21,22^ which could be released by cell-cell intercalation. Accordingly, we have shown that Akt downregulation is sufficient to increase clone splitting (EDFig. 7a,b).
2. The relative decrease of F-actin and tension in the loser-loser and winner-loser junctions (Fig. 4g). Overtime winner-loser junctions should get longer due to the relative lower tension, while winner-winner contacts should shrink due to higher tension. Consequently winner cells in contact with loser cells tend to get intermingled with loser cells, which can sometimes lead to cell-cell intercalation and separation of the winner cell from the rest of the winners.

The combination of the two forces increases the surface of contact between winners and losers and accelerates the elimination of loser cells by increasing the probability to induce loser apoptosis.

## Methods

### Fly stocks and clone induction

The following stocks were used in this study: *hs-flp22; tub<dmyc<Gal4; UAS-mcd8::GFP* ^4^, *hs-flp22; tub<dmyc<Gal4,ubi-Ecad::GFP; UAS-mcd8::RFP* (ubi-Ecad::GFP from ^31^, stock generated for this study), *UAS-diap1* (3^rd^, Bloomington), *UAS-fwe*^*loseA*^:*:HA* (on 3^rd^, ^12^), *UAS-fwe*^*loesA/B*^ *short hairpin RNAi* (3^rd^,^32^), *hs-flp22*; *act<y+<Gal4, UAS-GFP* (2^nd^, Bloomington), *hs-flp22*; *act<y+<Gal4, UAS-mcd8::RFP* (2^nd^,Bloomington); *hs-flp22; ubi-Cad::GFP, UAS-mRFP; act<y+<Gal4* (gift from Legoff L), *UAS-βgal* (2^nd^, Bloomington), *UASp-Ecad* (X,^33^), *UASt-sqhE20E21* (2^nd^, ^34^), *GMR-gal4, UAS-eiger* (2^nd^,^35^), *UAS-p35* (2^nd^, Bloomington), *zipper*^*2*^ (amorphic allele, Bloomington), *zipper*^*ebr*^ (hypomorphic allele, gfit from Baumann O), *dpp-gal4* (3^rd^, Bloomington), *hs-flp; UAS-fwe*^*loseB*^:*:HA; act<cd2<Gal4, UAS-GFP* (^12^), *act-gal4 switch 255* (2^nd^, ^36^), *FRT40A ubi-nlsGFP* (Bloomington), *FRT40A arm-βgal* (Bloomington), *FRT82B, arm-LacZ* (Bloomington), *FRT82B, RpS3, ubi-nlsGFP* (Minute mutant, Bloomington), *ubi-ptGPH::GFP* (2^nd^, ^20^), *UAS-myc* (3^rd^, Bloomington), *UAS-myc RNAi* (2^nd^, Bloomington), *hh-gal4,UAS-flp* (recombined 3^rd^,^37^ and Bloomington), *UAS-ptenRNAi* (3^rd^, Bloomington), *UAS-pi3kDN* (2^nd^, Bloomington), *UAS-pi3k*^*CAAX*^ (pi3kCA, X, Bloomington), *UAS-akt RNAi* (2^nd^, Bloomington), *UAS-akt* (2^nd^,Bloomington), Δ Δ[*dilps*_*1-*5_] (3^rd^, ^38^), sqh*-utABD::GFP* (2^nd^ or 3^rd^, ^39^) *UAS-Arp3 RNAi* (2^nd^, TRiP Bloomington), *UAS-Arp3::GFP* (2^nd^, Bloomington), *UAS-dia RNAi* (2^nd^, TRiP Bloomington), *UAS-dia::GFP* (3^rd^, ^40^), *FRT40A dia*^*5*^ (Bloomington, hypomorphic allele^40^), *endo-Ecad::GFP* (knock-in,^41^), *sqh-Sqh::GFP* (2^nd^, ^42^), *dachs-Dachs::GFP* (3^rd^, ^24^), *expanded-LacZ* (2^nd^, Bloomington), *fwe*^*loseA*^:*:mcherry* KI (generated for this study), *eng-Gal4, UAS-RFP* (2^nd^, Bloomington).

All heat shocks were performed in a 37°C waterbath using glass tubes. For notum live imaging, clones were generated using *hs-flp22; tub<dmyc<Gal4,ubi-Ecad::GFP; UAS-mcd8::RFP* line with a 35min heat shock or for 4hours (4 time 1 hours with 1 hour rest in between) for measuring the shape evolution of *tub-dmyc* patches surrounded by *wt* (EDFig.5d,e), 12min for *hs-flp;; act<y+<Gal4, UAS-mcd8::RFP* (crossed with ubi-Cad::GFP for *wt* in *wt* control) and *hs-flp; ubi-Cad::GFP, UAS-mRFP; act<y+<Gal4* (crossed with *UAS-fwe*^*loseA*^, *UAS-pi3kDN, UAS-pten RNAi, UAS-dia RNAi*) lines. For supercompetition assay in the wing disc (*wt* clones in *tub-dmyc*), larvae were heat shocked 20 min for movies, or 14min (Fig. 1e-g) or 12min (Fig. 2 to 4 and supplementary Figures). For *wt* clone in *wt* background (*hs-flp22*; *act<y+<Gal4, UAS-GFP*) we used 5min30sec heat shock for EDFig. 3a,b, and 5min for all other experiments. *UAS-fwe*^*loseB*^:*:HA; Act<cd2<Gal4* clones were induced with a 8min heat shock (EDFig.2). For *FRT40A ubi-nlsGFP/FRT40 arm-βgal* and *FRT40A dia5/FRT40 ubi-GFP* clones, larvae were heat shocked for 5min. We used 30min heat shocks to generate *wt* clones in *M-/+* background. Larvae were collected and dissected 24, 48 or 72h after heat shock. For starvation experiments, larvae were collected 24h or 48h after clone induction, washed in distilled water, and left on a humidified paper without food for 24h (48h for imaging pupae). Pupae were dissected and mounted as indicated in ^43^ 48 or 72h after clone induction.

Activation of progesterone sensitive gal4 (gal4 switch) was performed by using fly food mixed with RU486 at 1μg/ml or 50μg/ml. The full development was occurring in the hormone containing food (egg laying and larval development). *Act<cd2<Gal4* clones were induced with a 8min heat shock.

### Generation of Flower^loseA^::mcherry knock in

The Flower knock in fly was made by genomic engineering^41^. The genomic engineering by Huang is a two-step process consisting of ends-out gene targeting followed by phage integrase phi31-mediated DNA integration. A founder knock-out line was established with a genomic deletion of the *flower* locus at position 3L: 1’5816’737-15810028. The knock-in construct was composed of the deleted *flower* locus with a mCherry fusion after exon 5 (see EDFig.2a, specific for *loseA* isoform). The knock-in construct was done by site directed mutagenesis to remove the stop codon and add a restriction site, used to insert mCherry after exon 5. The knock-out of *flower* and the knock-in *Flower*^*loseA*^:*:mcherry* were tested by PCR and sequencing.

*Vectors used for generating the FlowerloseA::mcherry:*

pGX-attP: Knock-out vector

pGEM-T: Used for the site directed mutagenesis

pGEattB^GMR^: Knock-in vector

*Primers:*

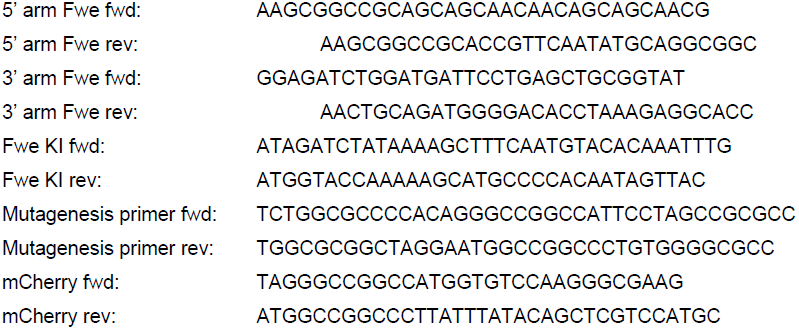

### Immunohistochemistry and image acquisition

Immunostainings of the wing discs were performed with standard formaldehyde fixation and permeabilisation/washes in PBT 0.4% Triton. The following antibodies/markers were used: rat anti E-cad (DCAD2 concentrate, DSHB, 1/50), chicken anti GFP (1/200, Abcam), guinea-pig and rabbit anti Dia (1/50, gift from J. Grosshans, ^44^), rabbit anti phospho-Akt (1/50, Cell Signalling), guinea-pig anti Myc (1/500, gift from G. Morata), phalloidin Alexa 546 (1/50, Life technologies), mouse anti Ptc (1/50, DSHB), rabbit anti β-gal (1/200, Cappel), rat anti HA (1/100, Roche). Dia and phospho-Akt stainings were amplified using biotinylated secondary antibodies and streptavidin coupled to Cy3 or Cy5. Tunnel stainings were performed using the Roche kit.

Dissected wing disc were mounted in Vectashield (Vectorlab) and imaged on a Leica confocal SP2 using a 63X water immersion objective N.A. 1.3 or a Leica confocal SP8 using a 63X oil objective N.A. 1.4. Images shown are maximal z-projection containing the adherens junction plane.

Adult eye pictures were taken on a Leica MZFLIII dissecting scope with a Nikon DXM1200 colour camera at the same magnification. Eye area was measured on Fiji ^45^.

### Pupal notum and wing disc live imaging, death probability calculation

Pupae were collected 48 or 72h after clone induction and dissected 18 to 20h after pupae formation (APF). Pupae were prepared as indicated in ^43^ and imaged on a confocal spinning disc microscope (Till photonics) with a 40X oil objective (N.A. 1.35) or a point scanning confocal microscope Leica SP8 with a 63X objective (N.A. 1.4). Large field views of the tissue were obtained by doing tile imaging (6 to 12 tiled positions). Z-stacks (1 μm/slice) were recorded every 5min using autofocus at every cycle and every tiled position using E-cad::GFP signal. Movies were performed in the nota close to the scutellum region in the vicinity of the aDT and pDT macrochaetes. Movies shown are cropped from larger field of views after maximum projections, correction for bleach (using Fiji) and correction for tissue drift (Stack reg plugin, Fiji).

Every cell delamination event in the clones was tracked over 10h. We excluded cells in the midline where spontaneous delaminations are occurring^46^. The proportion of apical perimeter shared with winners (sum of the winner-loser junction lengths over the total apical perimeter) was measured 1 hour prior to delamination at the junction plane (E-cad::GFP signal or actin belt visualised with utABD::GFP) using imageJ. We then measured the proportion of apical perimeter shared with winners for all the cell in the clones at time 0 in every movie (excluding cells in the midline) manually or using CellPacking analyser for skeletonisation^47^ and a home-made macro (Igor Pro Software, Wavemetrics). Death probabilities were then obtained by dividing the number of delaminating cells by the total number of loser cells in each shared perimeter category. Note that similar correlations were obtained by using the number of junctions shared with winners divided by total number of junctions (not shown).

*Ex-vivo* culture of wing-disc was performed using clone8 media, as indicated in ^21^, using a point scanning confocal microscope Leica SP8 with a 63X objective (N.A. 1.4), 36h after clone induction (*wt* in *tub-dmyc*). Discs showing rapid drop down of cell division were excluded from the analysis. Removal of the signal from the peripodal cells and selection of the signal from the junction plane were performed by using a home made Matlab macro for selective plane projection (inspired by ^21^). For every x-y pixel, the z-plane with the maximal E-cad::GFP intensity (calculated by summing pixel intensity on a 50 by 50 px square at every plane) was kept and also used to retrieve RFP signal in the same plane. The measurement of death probability was performed like in the notum. Imaging of *fwe*^*loseA*^:*:mcherry* KI was also performed in *ex-vivo* wing disc using the same projection procedure.

Junction remodelling were manually counted at the interface of the clones. Each event leading to the disappearance of a junction between two RFP positive cells was counted. Remodelling events were only counted if the new topology was maintained until the end of the movie (10h, Fig. 2f and EDFig.5c). The total number of remodelling events was then divided by the total number of junctions analysed. For EDFig.5a, we counted every remodelling event occurring over 10h for single junctions, and calculate the proportion of junctions undergoing a single remodelling event, and the probability to undergo additional remodelling event (after a first remodelling event). All winner-winner junctions tracked were sharing one vertex with a loser cell, while loser-loser junctions tracked were also sharing one vertex with a winner cell. All winner-loser junctions were tracked.

### Image quantification

#### Clone fragmentation, clone size and clone compactness

Clones were counted as fragmented when GFP positive cells were separated by a single GFP negative cell in the apical area (using E-cad or phalloidin staining) 48h after clone induction in the wing pouch. Discs showing too high clone density were excluded (>20 clones/wing pouch). This technics accurately evaluates the number of clones that have recently split. The probability to obtain false positives is expected to be low as the probability to observe two independent clones separated by a single cell requires the initial recombination to occur in two cells separated by a single cell, as well as the absence of division of the cell in between for the subsequent 48h (on average, cells should divide more than 3 times ^48^). Accordingly, we did not observe a lower number of fragmentation when it was evaluated using twin clones (EDFig.4a,b) where 2xβ-gal positive clones are counted as fragmented when cells are separated by single negative cell, and when the sister clone (2xGFP) forms a single continuous group. However, we slightly underestimate the total number of fragmented clones as the proportion of all the split 2xβ-gal clones was close to 12% (including groups separated by more than 1 cell) (**EDFig.4a,b**).

Clone size was measured on 9μm wide maximum projection containing the apical junction plane in the wing pouch. GFP positive patches were automatically recognized and segmented using a home-made Fiji macro, and used to measure the average clone area, the density of clone ([number of cloned / area of the wing pouch] x 10,000 = number of clones per 10,000μm^2^), and the total surface of the wing pouch covered by GFP positive cells (total clone area/area of the wing pouch).

Clone compactness is defined as followed:

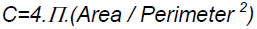

Clone compactness was measured for every clone in the notum by drawing their contour using Fiji at t0 and 10 hours later. The fold change was calculated as followed: (C_10h_-C_0h_)/C_0h_.

#### Death induction at compartment boundary

Measurement of death induction at the AP compartment boundary was performed by counting the number of tunnel positive cells in the wing pouch in the posterior compartment (control compartment) and in a ROI in the anterior compartment (a band of 3 cells width along Antero-Posterior compartment boundary). Compartment boundary was detected with Patched staining. Z-projection of the three different genotypes (1. *dpp-gal4/UASfweloseA::HA,* 2. *zip*^*2*^*/zip*^*ebr*^, 3. *zip*^*2*^*/zip*^*ebr*^; *dpp-gal4/UASfweloseA::HA*) were randomized by assigning random file name and tunnel cell counting was performed blindly. The expected random distribution of dead cells in the ROI was calculated by taking the average ratio of surface of the ROI over the total surface of the wing pouch for 10 wing discs. *zip*^*2*^*/zip*^*ebr*^ larvae were sorted by using a fluorescent balancer chromosome (*Cyo, act-GFP*, Bloomington).

#### Intensity measurements

*fwe*^*loseA*^:*:mcherry* intensity measurements were performed in ex-vivo cultured wing disc on selective plane projections (see above). For each cell in the loser clones, we measured the membrane intensity using Fiji (6px width line) divided by the average membrane intensity of all the cells measured in the same disc, and measured the percentage of perimeter shared with winner cells.

PIP3 intensity measurements were performed on maximum projections of cell apical area (4 μm) in living notum expressing ubi-ptGPH::GFP. Cytoplasmic signal was removed using two subsequent background substractions on Fiji (Rolling Ball radius 200px and 5px). Junction signal is the mean intensity in a 6 px width line (1px=0.148 μm). Measurements of utABD::GFP junction intensity were performed similarly on a 6px width line after a single background subtraction (rolling ball radius 200px).

Measurement of line intensity profile in the wing discs (EDFig.10) were performed by using a 100px width line in the dorsal compartment parallel to the dorso-ventral compartment boundary (1px=0.267μm) on maximal z-projections using Fiji. Intensity profiles were measured for Patched (Ptc) and utABD::GFP or Dia. Position 0 was determined by detecting the minimum of the derivative of Ptc intensity profile (calculated on Igor Pro Software after smoothing of the profile, 300px averaging window), which corresponds to the boundary between the anterior and the posterior compartments. The distance from the most anterior side of the profile to the AP boundary was normalized to 1. Each intensity profile was divided by its mean. The average intensity profile was then calculated by averaging the normalized intensity values obtained for each disc at the same relative distance to AP boundary.

#### FRAP

utABD::GFP has been previously used to assess actin dynamics^39,49^. *utABD::GFP* FRAP experiments were performed in *ex vivo* cultured wing disc in DMSO (0.2%, control) and after treatment with Jasplakinolide (2μM, life technologies) using a confocal spinning disc microscope (Till Photonics) and a 60x oil objective (N.A.: 1.35). Bleaching was performed using 100% of the power of a 488 solid state laser in diffraction limited ROI after acquiring 3 time points (1 frame/0.5sec). Recovery was recorded on 30sec. Intensity recoveries were obtained by measuring the mean intensity in a 15x15px ROI on Fiji (1px=0.10μm) containing the bleached region. Each curve was normalized by the intensity profile of a neighbouring control region (20x20px) to correct for bleaching due to imaging. For each normalized curve, we subtracted the intensity at t0 (post bleach) and then divided by the average intensity of the 3 first points (pre-bleach). The averages of all the recovery curves are then shown. FRAP in the notum were performed similarly 48h after clone induction (*wt* losers in *tub-dmyc*). Characteristic times of recovery were calculated by fitting the normalized curves with Igor Pro software using the following equation:

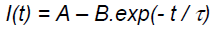

Where t is the time (in sec), τ is the characteristic time of recovery, A the mobile fraction and (A-B) the initial intensity after bleaching. For comparison between winner-winner and winner-loser junction in the same cell, we performed simultaneous bleaching and recovery recording in two identical ROI (one in a winner-winner junction, one in a winner-loser junction of the same cell). Each recovery curve was normalized and fit as mentioned above. We then calculated the fold change of characteristic time of recovery for each cell:

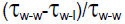

*endo-Ecad::GFP* FRAP experiments were performed similarly using a Leica SP8 point scanning confocal (63X oil immersion objective, N.A. 1.4), using an Argon laser for imaging and bleaching (488nm). Recovery was assessed over 200sec, 1 frame/sec. The average recovery curves were obtained like *utABD::GFP* FRAP experiments.

### Laser ablation

Junction laser ablations were performed in the notum using a two-photon infrared laser on a 2-photon microscope (Two-photon Fluoview 1000 Olympus, Center for Microscopy and Image Analysis, University of Zurich) using a 25X water objective (N.A. 1.05). Imaging and ablation were performed with 950nm wavelength, 1.3% power for imaging, 30% power and 300ms exposition for the ablation scanning along a line perpendicular to the junction. Relaxation of vertices (visualised with E-cad::GFP) was tracked for at least 10sec (1 frame/0.594sec, 1 frame/0.891sec for Fig. 4f “starved”). Vertices position was then tracked using CellTrack software^50^ and the evolution of vertices distance over time was fitted with an exponential function (A_0_+A_1_.exp(A_2_.t)) on the ten first time point following ablation using Igor software. The V_max_ were the derivative of the exponential fit at t0.

### Statistics

All the error bars shown in the figures are Standard Errors of the Mean (s.e.m.) or 95% confidence interval for death probabilities. Statistical tests are Fisher exact tests (two sided) for comparison of proportions or Mann-Whitney non-parametric tests for all the other experiments, except in EDFig.8f where we used a one-sample t-test with 0 as reference value (normality tested with a Shapiro-Wilk test). Difference in variance was always below 2. No statistical methods were used to set sample size. Experiments were not randomized, and we did not analyse experiments blindly, except for the result of Fig. 1h. Every experiment was performed at least 3 independent times. The same control is used for all the quantification of clone splitting as it is based on 3 independent experiments and give reproducible values (for example compare **Fig. 2d**, *wt in wt* and **EDFig.4a,b**).

## Extended data figure legends

### Extended data Figure 1: Loser elimination also correlates with the shared perimeter with winners in the wing disc

**a,** Left: Selective plane Z-projection of an ex-vivo cultured wing discs expressing ubi-Ecad::GFP with loser clones (RFP, purple, *wt* in *tub-myc*) 36h after clone induction. 1,2,3 correspond to the clones shown on the right. Right: snapshots of loser clones showing the delamination of loser cells and one event of clone splitting preceding cell elimination (3). Scale bars=5μm. **b**, Distribution of the proportion of junctional perimeter of loser cells shared with winners in eliminated cells 1 hour prior to delamination (top, 0=cell in the center of the clone, 1=isolated cell surrounded by winners) and in all the loser cells at t0 (bottom). **c,** Probability of loser cell elimination for a given surface of contact shared with winners in *wt* loser cells in *tub-dmyc*. Statistical tests are Fisher-exact tests performed with the point 0-0.25 (**-**: p>0.05, *:p<0.05, **:p<10^−2^, ***: p<10^−3^, ****: p<10^−4^). Error bars are 95% confidence interval.

### Extended data Figure 2: Contact dependent death is triggered downstream of *flower*

The transmembrane protein Flower is a central regulator of competition^12^. *fwe*^*lose*^ isoforms (*loseA* and *loseB*) are induced downstream of several competition contexts and their expression is necessary for loser elimination and sufficient to drive cell elimination when contacting *wt* cells^12^. The contact dependent communication could occur upstream of *fwe* (for instance by modifying the levels of induction of *fwe*^*lose*^) or downstream of *fwe* induction. Several evidences indicate that it occurs downstream of *fwe* induction. First, cell death also correlated with shared apical perimeter in clones homogenously expressing *fwe*^*loseA*^ (Fig. 1c, red curve). Secondly, using a knock-in fusion *fwe*^*loseA*^:*:mcherry* (**EDFig.2a**), we could show that *fwe*^*lose*^ induction did not correlate with the surface of contact shared with winners (**EDFig.2b,c**) as previously suggested by *in situ* for *fwe*^*lose*^ ^12^. Finally, the probability of elimination of clones overexpressing *fwe*^*lose*^ is proportional to the relative differences in *fwe*^*lose*^ levels inside and outside the clones (**EDFig.2d,e**). Altogether, this suggested a model where cells can compute the relative differences of *fwe*^*lose*^ levels with all their neighbours through an unknown molecular mechanism.

**a,** Schematic of the modified *fwe* locus (left) and the resulting mRNA of the 3 isoforms (right). Orange rectangles are exons. The 5’ and 3’ UTR are shown in purple. Exon 5 is specific to each isoform. The red box shows the localisation of the mCherry tag ad at the end of the exon 5 of *fwe*^*loseA*^. Note that the vector backbone was conserved in the *knock in* line (white, AmpR).**b**, Two examples of selective plane Z-projection of ex-vivo cultured wing discs expressing *fwe*^*loseA*^:*:mcherry KI* in *wt* clone in *tub-dmyc* background (purple) 36h after clone induction. Clone contour is shown in purple (right). 12/12 discs. Scale bars=10μm. The intensity profile of the white dot line is shown below. Bottom right panel shows a lateral view of *fwe*^*loseA*^:*:mcherry* and its accumulation in the apico-lateral region. **c**, Scatter plot of *fwe*^*loseA*^:*:mcherry* membrane intensity in loser cells in wing disc (*wt* in *tub-myc,* y axis) against the percentage of perimeter shared with winner (x axis). One dot=one cell. Pearson Correlation Coefficient=-0.24. **d**, *UAS-fwe*^*loseB*^:*:HA* clones (GFP) in wing discs 72h after clone induction (ACI) with different concentrations of RU486 in the food media. *fwe*^*loseB*^ expression in clones is the sum of act-G4 flip out driver (constant) and the hormone sensitive Gal4 (Gal4 switch, expression proportional to the RU486 concentration) while *fwe*^*loseB*^ is only driven by the Gal4 switch outside the clone. GFP panels were acquired with the same parameters and are shown with the same contrasts. Scale bars = 100μm. **e**, Average percentage of the wing pouch surface covered by clones (left, purple histogram) and average GFP intensity ratio inside/outside clones (right, green curve, log scale) 72h ACI. Error bars are s.e.m.. Statistical tests are Mann-Whitney tests performed for wing pouch coverage. **f**, *UAS-fwe*^*loseB*^:*:HA* clones (GFP) in wing discs 72h after clone induction (ACI) in a control (left, *UAS-lacZ*) or upon overexpression of active MRLC (*UAS-sqhE20E21*, right). Scale bars = 100μm. **g**, Average percentage of the wing pouch surface covered by clones 72h ACI. Error bars are s.e.m.. Statistical test is a Mann-Whitney test.

### Extended data Figure 3: E-cad and active MRLC rescue losers only through the change of winner/loser surface of contact

**a,** Supercompetition assay in the wing disc 24h, 48h and 72h after clone induction (purple: loser cells) in normal competition (loser cells overexpressing β-gal), upon limitation of cell mixing (*UAS-ecad* and *UAS-sqhE20E21,* a constitutively active MRLC) and in *wt* clones in *wt* background (no Competition). Left: schematic of the expected effect on clone shape, black line thickness is the strength of cell-cell adhesion, red lines show actomyosin network. Middle: example of wing discs at different time after clone induction, scale bars = 100μm. Right: close up views of clones overexpressing E-cad (top) and active MRLC (bottom). Active MRLC induces apical actin accumulation and partial apical constriction. Scale bars = 5μm. **b,c,** Density of loser clones (**b**) and averaged loser clone size (**c**) at 24,48 and 72h after clone induction (ACI) in the wing pouch. Error bars are s.e.m.; n are the number of wing discs (**b**) or the number of clones (**c**). Statistical tests are Mann-Whitney tests performed with the control competition (green, *UAS-βgal* in *tub-dmyc*) or with control without competition (orange, *wt* in *wt*) **-**: p>0.05, *:p<0.05, **:p<10^−2^, ***: p<10^−3^, ****: p<10^−4^. Note that we did not find significant differences of clone size at 72h as the few clones remaining for the control competition were at the periphery of the wing disc where competition is less effective. **d,** E-cad and active MRLC do not have a cell autonomous effect on growth. Left: Wing discs showing *wt* clones (*act<y<gal4, UAS-mcd8::GFP*, purple) overexpressing *β-gal*, *E-cad* or *sqhE20E21* (active MRLC) 48h ACI. The insets show the clones marked with an asterisk. Scale bars=100μm. Right: Average clone surface, n=number of clones, error bars are s.e.m.. **-**: p>0.05, Mann Whitney tests. **e,** E-cad and active MRLC do not prevent genetically induced apoptosis. Left: Adult eye of a *wt* fly (*oregonR*), and flies with abnormal eye morphology due to induction of JNK dependent death in the eyes (eye specific gal4, GMR-gal4 and UAS-eiger, the fly ortholog of TNF^35,51^) expressing *β-gal* (control), *diap1* (apoptosis inhibition), *E-cad* or *sqhE20E21* (active MRLC). Right: Averaged eye surface in pixel (a.u.: arbitrary units). **f,** E-cad and active MRLC do not modify loser death probability for a given surface of contact with winner. Probability of loser cell elimination in the pupal notum for a given surface of contact shared with winners in Myc-dependent competition (purple, from Fig.1d), in *wt* cells in *wt* background (control, dashed green, from Fig. 1d), or in losers overexpressing E-cad (dashed red) or active MRLC (*sqh-E20E21,* dashed pink). Statistical tests are Fisher-exact tests performed with Myc-dependent competition (purple) (**-**: p>0.05). Error bars are 95% confidence intervals.

### Extended data Figure 4: Clone fragmentation does not correlate with clone size

**a,** Twin clones 48h ACI marked with 2 copies of GFP (green) and absence of β-gal or 2 copies of β-gal (red) and absence of GFP (*FRT40A ubi-nlsGFP/FRT40A bcat-βgal*). Left: Non-fragmented clones. Middle and right show fragmented clones (the GFP sibling clone is used as a reference) with clone cells separated by a single cell (middle) or more than one cell (right). **b,** Proportion of fragmented clones 48h ACI in *wt* GFP clones in *wt* background (blue, from Fig. 2d) quantified with the one cell distance criteria. Same quantification in *FRT40A ubi-nlsGFP/FRT40A bcat-βgal* where 2x*βgal* clones were counted as split when clone cells were separated by a single cell and were associated with a continuous group of sibling *2xGFP* cells. This quantification showed no differences with the *wt* GFP clones in *wt* background, demonstrating that our method does not produce false positives. However, it slightly underestimates the total number of fragmented clones (compare with “all” where every split 2x*βgal* clones is counted). n= number of clones. Statistical tests are Fisher-exact tests performed with *wt* GFP clones in *wt* background (blue). **c**, Representative wing discs 48h ACI in control (*wt* in *wt*) and in supercompetition assay with loser cells expressing *β-gal*, *UAS-ecad*, *UAS-sqhE20E21*, *UAS-diap1* (an endogenous apoptosis inhibitor) or *UAS-p35* (a bacterial Caspase 3 inhibitor) and *fwe*^*lose*^ RNAi; or after induction of winner clones (*UAS-p35*,*UAS-myc* in *wt, p35* is necessary to block the cell autonomous death induced by high Myc overexpression^52^, and *wt* in *M-/+* where *wt* clones have no GFP). White arrowheads show fragmented clones. Insets show close up view of representative clones. Scale bars = 100μm. **d**, Scatter plot showing the proportion of fragmented clone (y axis) against the average size of clone (x axis) 48h after clone induction for all the different genotype used in this study (see legend). One dot=one fragmentation assay. There is no correlation between clone size and clone splitting. Pearson Correlation Coefficient=0.14. Note that also without the outlayer point (*UAS-pten RNAi* in *wt*) the correlation is close to 0, corr.coef.=-0.036.

### Extended data Figure 5: Clone fragmentation is driven by winner-loser mixing

**a,** Left: schematic showing loser cells (*wt*, purple) and winner cells (*tub-dmyc*, green). Orange junctions are junctions shared by a winner and a loser cell (winner-loser junctions). Dark green junctions are the winner-winner junctions (sharing one vertex with a loser cell) and dark purple are the loser-loser junctions (sharing one vertex with a winner cell) used for the analysis. Right: proportion of junction undergoing a single remodelling event over 10h in the notum. p-values=Fisher exact tests. **b**, Probability to undergo additional junction remodelling after a first remodelling event. n=number of junctions. p-values=Fisher exact test. This suggests that winner-loser junctions undergoing a first remodelling events have a higher probability to revert back to the initial topology. **c**, Left: Snapshots of *wt* cells and *tub-dmyc* cells in the notum (no clone) at t0. Purple junctions disappear after 10h while green junctions remain unchanged (see Video S12). Scale bars=10 μm. Right: proportion of junction disappearing after 10h. p-values=Fisher exact tests. **d**, Examples of clone in the notum at t0 and 10 hours later for various genotypes. E-cad::GFP is in green and UAS-RFP in purple. The white dashed lines show clone contours. Scale bars=10 μm. **e**, Fold change of clone compactness after 10h in the notum (see Methods). One dot=one clone. The bars are averages. p-value=Mann-Whitney tests (**-**: p>0.05, *:p<0.05, **:p<10^−2^, ***: p<10^−3^, ****: p<10^−4^).

### Extended data Figure 6: Differential PIP3 drives clone splitting

**a,b,** Z-projection of tGPH (PIP3, green and pseudocolour) in the pupal notum in clones (RFP, purple) overexpressing a dominant negative of PI3K (*UAS-pi3kDN*, **a**) or upon downregulation of PTEN (*UAS-pten RNAi*, **b**). White dashed lines show clone boundaries. Scale bars = 10μm. **c**, Z-projection of a phospho-Akt staining (green) in wing disc overexpressing *fwe*^*loseA*^:*:HA* in the posterior compartment (purple, *eng-G4,* 16/16 discs). Scale bar = 100μm. **d,** phosphor-Akt in *wt* clones (no GFP) surrounded by *M-/+* cells (21 discs). White arrows point to some *wt* clones. Scale bar = 100μm. **e**, Z-projections of phospho-Akt (green), GFP (magenta) in wing discs after removal of one additional copy of Myc in the posterior compartment and 24h of starvation (*hh-gal4*, *UAS-flp* x *tub>dmyc>gal4, UAS-GFP,* 10/10 discs). Scale bar=100μm. **f,** Z-projection of tGPH (PIP3) in loser clones (supercompetition assay, purple are losers) in the pupal notum after 48h of starvation. Scale bars = 10μm. **g,** Quantification of the mean junction membrane intensity of tGPH in winner-winner, loser-loser and winner-loser junctions in the notum. n=number of junctions. Error bars=s.e.m. p-values: Mann-Whitney tests. **h,** Wing discs with loser cells (supercompetition assay) 48h ACI after 24h of starvation, or upon removal of one copy of *Dilps 1 to 5* (Drosophila Insulin Like Peptides). Insets show representative clones. Scale bars=100μm. **i,** Proportion of fragmented clones. n= number of clones. Statistical tests are Fisher-exact tests performed with *wt* (blue) or control competition (green), **-**: p>0.05, *:p<0.05, **:p<10^−2^. *wt* in *wt* and *wt* in *tub-dmyc* come from Fig. 2d. This result suggests that loser and winner have differential ability to process and respond to extracellular insulin. **j**, Wing discs showing *UAS-βgal* clones (loser, purple) in *tub-dmyc* background 72h after clone induction (ACI) with or without a 24h starvation period. Scale bars = 100 μm. **k**, Average density of loser clones (left) and average percentage of the wing pouch covered with GFP positive cells (right) 72h ACI. Error bars are s.e.m.. p= Mann Whitney tests.

### Extended data Figure 7: Akt is not sufficient to explain winner-loser mixing and E-cad, MyoII and Dachs do not show visible defects in loser clones

**a**, Wing discs showing clones upon downregulation of Akt (*UAS-akt RNAi*, left) or upregulation of Akt (*UAS-akt*, right) 48h ACI. Insets show close up view of a representative clone in each condition. Scale bars = 100μm. **b,** Proportion of fragmented clones. *wt in wt* comes from Fig. 2d. *UAS-pi3kDN in wt* and *UAS-pten RNAi in wt* come from Fig. 3d,e. n=number of clones. Statistical tests are Fisher-exact tests performed with *wt* (blue) or as indicated by the dashed lines, **-**: p>0.05, *:p<0.05, **:p<10^−2^, ***: p<10^−3^, ****: p<10^−4^. **c,** Z-projection of endo-Ecad::GFP (knock-in line) in the pupal notum. Loser clones are marked with RFP (*wt* in *tub-dmyc* background, 3 nota, 30 clones). White line marks clone contour. Scale bar=10μm. **d**, Average normalized intensity recovery curves of endo-Ecad::GFP after photobleaching in loser-loser junctions (*wt in tub-dmyc*, purple) and in winner-winner junctions (*tub-dmyc*) in the notum. Error bars=s.e.m.. **e,** z-projection of MRLC::GFP (endogenous promoter, *spaghetti-squashed, sqh*) in the pupal notum. Loser clones are marked with RFP (*wt* in *tub-dmyc* background, 5 nota, 30 clones). White line marks clone contour. Scale bar=10μm. Note that utABD::GFP (as shown in Fig.4a) is under the control of the same promoter, thus the actin reduction in losers is not due to a reduction of *sqh* promoter activity. **f,** z-projection of Dachs::GFP (endogenous promoter) in the pupal notum. Loser clones are marked with RFP (*wt* in *tub-dmyc* background, 2 nota, 17 clones). White line marks clone contour. Scale bar=10μm.

### Extended data Figure 8: F-actin turnover/polymerisation rate is reduced in loser junctions

utABD::GFP (utrophin Actin Binding Domain) has been previously used to assess actin dynamics^39,49^. This is further demonstrated by the experiment described in **a** and **b**. **a**, *ex-vivo* cultured wing discs in control media (DMSO 0.2%) or in media containing 2μM of Jasplakilonide, an inhibitor or actin turnover^53^. The white rectangles are the bleached zones. White dash lines are the one used for the kymographs shown on the right. t=0s (white dashed line on kymographs) is the time of bleaching. Scale bars = 5μm. **b**, Averaged normalized recovery curves of utABD::GFP intensity after photobleaching in control and Jasplakinolide treated wing discs. Error bars are s.e.m.. **c**, Averaged utABD::GFP normalized intensity recovery curves in loser-loser junctions (purple), winner-winner junctions (green) and winner-loser junctions (orange) after photobleaching (*wt* losers in *tub-dmyc*) in the notum. Error bars are s.e.m.. **d,** Distribution of the characteristic times of utABD::GFP intensity recovery in winner-winner, loser-loser and winner-loser junctions. p-values=Mann-Whitney tests. **e,** Top, schematic of the FRAP experiments, two ROIs are bleached simultaneously in the same winner cell (*tub-dmyc*) sharing contacts with a loser cell (*wt*). Grey square, winner-loser bleached junction, black square, winner-winner bleached junction. Bottom, confocal image in the pupal notum 48h ACI of utABD::GFP in a supercompetition assay (purple cells: losers). Scale bar=5μm. Squares show the simultaneously bleached regions (1: winner-winner junction, 2: winner-loser junction), the white dashed line is the line used for the kymograph shown on the right. **f,** distribution of the fold change of the characteristic time of intensity recovery in the winner-winner junction compared to a winner-loser junction of the same cell ((τ_w/w_-τ_w/l_)/τ_w/w_), one dot=one cell. The bar is the average. The statistical test is a one-sample t-test with 0 as reference value.

### Extended data Figure 9: Filamentous actin defects are necessary and sufficient to drive clone fragmentation

**a**, Z-projection of phalloidin (green) and GFP (magenta) in a wing disc containing *wt* clones (no GFP) in *M-/+* background. Top inset shows phalloidin signal for two *wt* clones (white lines). Right: close up views of cell shape in two *wt* clones**. b,** Representative wing discs 48h ACI upon silencing of Arp3 (*arp3 RNAi*) in *wt* background, in supercompetition assay with loser cells expressing Arp3::GFP, silencing of Diaphanous (a formin, *dia RNAi*) in *wt* background, and in supercompetition assay with loser cells expressing Dia::GFP. Discs correspond to experiments quantified in Fig. 4d, control *wt* and control supercompetition assay are the same as in Fig. 2d. White arrowheads show fragmented clones. Insets show close up view of a representative clone in each condition. Scale bars = 100μm. **c,** Wing disc 48h ACI. The clones without GFP are homozygous mutant for *diaphanous* (dia^5^, hypomorphic allele^40^), the sibling *wt* clones have two copies of GFP. Four close up views of fragmented mutant clones are shown on the right. The proportion of fragmented clone is 39.3% (n=132 clones), counting every fragmented clone (including patches separated by more than one cell). Scale bar=100μm. **d,** Top: representative wing discs during supercompetition 72 ACI in loser cells (purple, GFP) overexpressing β-gal (control) or Dia::GFP. Scale bar=100μm. Bottom: Quantification of the mean loser clone density and the average proportion of the wing pouch surface covered by loser clones. *:p<0.05, **:p<10^−2^. Mann-Whitney test. Error bars are s.e.m.. **e,** *expanded* level of expression (*exp-lacZ*), a downstream target of Yki/YAP, in two representative examples of clones overexpressing Dia::GFP (*hs-flp22, act<y<gal4; UAS-GFP x UAS-dia::GFP*) 72h ACI (same result for 31 clones, 16 wing discs). White lines show the contour of the clones. Scale bars=5μm.

### Extended data Figure 10: PIP3 acts upstream of actin defects

Starvation was sufficient to abolish differences in F-actin between compartment expressing different levels of Myc (**EDFig.10a**), while overexpression of *fwe*^*loseA*^ in the posterior compartment did not modify F-actin (**EDFig.10b**). Moreover, PIP3 downregulation in a full compartment was sufficient to downregulate actin (**EDFig.10c**) and junctional Diaphanous (**Ext Fig. 10d**). Finally, overexpression of Dia significantly reduced the number of fragmented clones upon downgregulation of PIP3, while knocking down Dia and increasing PIP3 impaired the rescue of loser fragmentation (**EDFig.10e,f** p=0.017 and 0.003 respectively). Altogether, we concluded that actin defects are driven by the modulation of PIP3 in loser cells.

**a,** Z-projections of utABD::GFP (green, F-actin), RFP (magenta) in wing discs after removal of one additional copy of Myc in the posterior compartment (*hh-gal4*, *UAS-flp* x *tub>dmyc>gal4, UAS-RFP*, left, 20/20 discs) and after 24h of starvation (right, 9/10 discs). Scale bars=100μm. **b**, Z-projection of utABD::GFP (green, F-actin) in a wing disc overexpressing *fwe*^*loseA*^:*:HA* in the posterior compartment (purple, *eng-G4,* 20/20 discs). Scale bar = 100μm. **c**, utABD::GFP and Phalloidin stainings in control wing discs (*hh-gal4* alone) and upon reduction of PIP3 in the posterior compartment using *hh-gal4* and *UAS-pi3kDN* (see top scheme, A: Anterior, P: Posterior, D: Dorsal, V: Ventral). The posterior compartment is at the right side of the Patched (Ptc) stripe marked in blue. Scale bars=100μm. Bottom: Averaged normalized intensity line profiles along Antero-posterior axis for utABD::GFP (green and yellow) and Ptc (blue). Position 0 corresponds to the maximum inflexion of Ptc intensity peak (right side of the stripe). n=number of wing discs. Error bars are s.e.m.. **d,** Diaphanous staining in control (*hh-gal4* alone) and upon reduction of PIP3 in the posterior compartment using *hh-gal4* and *UAS-pi3kDN*. Scale bars=100μm. Bottom: averaged normalised intensity line profile taken along Antero-posterior axis for Dia (green and yellow) and Ptc (blue). n=number of wing discs. Error bars are s.e.m.. **e,** Wing discs 48h ACI showing clones (*hs-flp22; act<y<gal4; UAS-GFP* purple) in *wt* background overexpressing a dominant negative form of PI3K *(UAS-pi3kDN*) and *dia::GFP*, or loser clones in *tub-dmyc* background overexpressing a constitutively active form of PI3K (*pi3kCA*) and *dia RNAi*. The white arrowhead shows a fragmented clone. Insets show close up views of a representative clone in each condition. Scale bars = 100μm. **f,** Proportion of fragmented clones. Blue bars are *act<y<Gal4* clones in *wt* background, green bars are loser clones in *tub-dmyc* background. *wt in wt* and *UAS-βgal* in *tub-dmyc* (supercompetition assay) come from Fig. 2d. *UAS-pi3kDN in wt* and *pi3kCA in tub-dmyc* come from Fig. 3d,e. n=number of clones. Statistical tests are Fisher-exact tests performed with *wt in wt* (blue) or *UAS-βgal* in *tub-dmyc* (green) or as indicated by the black dashed lines, **-**: p>0.05, *:p<0.05, **:p<10^−2^, ***: p<10^−3^, ****: p<10^−4^.

